# Meta-Prism 2.0: Enabling algorithm for ultra-fast, accurate and memory-efficient search among millions of microbial community samples

**DOI:** 10.1101/2020.11.17.387811

**Authors:** Kai Kang, Hui Chong, Kang Ning

## Abstract

**Motivation:** Microbial community samples and sequencing data have been accumulated at a speed faster than ever, with tens of thousands of samples been sequenced each year. Mining such a huge amount of multi-source heterogeneous data is becoming more and more difficult. Among several sample mining bottlenecks, efficient and accurate search of samples is one of the most prominent: Faced with millions of samples in the data repository, traditional sample comparison and search approaches fall short in speed and accuracy.

**Results:** Here we proposed Meta-Prism 2.0, a microbial community sample search method based on smart pair-wise sample comparison, which pushed the time and memory efficiency to a new limit, without the compromise of accuracy. Based on memory-saving data structure, time-saving instruction pipeline, and boost scheme optimization, Meta-Prism 2.0 has enabled ultra-fast, accurate and memory-efficient search among millions of samples. Meta-Prism 2.0 has been put to test on several datasets, with largest containing one million samples. Results have shown that firstly, as a distance-based method, Meta-Prism 2.0 is not only faster than other distance-based methods, but also faster than unsupervised methods. Its 0.00001s per sample pair search speed, as well as 8GB memory needs for searching against one million samples, have enabled it to be the most efficient method for sample comparison. Additionally, Meta-Prism 2.0 could achieve the comparison accuracy and search precision that are comparable or better than other contemporary methods. Thirdly, Meta-Prism 2.0 can precisely identify the original biome for samples, thus enabling sample source tracking.

**Conclusion:** In summary, Meta-Prism 2.0 can perform accurate searches among millions of samples with very low memory cost and fast speed, enabling knowledge discovery from samples at a massive scale. It has changed the traditional resource-intensive sample comparison and search scheme to a cheap and effective procedure, which could be conducted by researchers everyday even on a laptop, for insightful sample search and knowledge discovery. Meta-Prism 2.0 could be accessed at: https://github.com/HUST-NingKang-Lab/Meta-Prism-2.0.

## Introduction

Microbial communities have asserted great influences in healthcare, environment and industry[1–4]. As such, an increasing amount of projects have been conducted on microbial communities around the world, such as those from the “Human Microbiome Project”[1, 2] and the “Earth Microbiome Project”[3, 4]. These massive amount of samples have already discovered knowledge about microbial community and their effects on environment and human health[5, 6], providing opportunity to study the hidden evolution and ecology patterns among microbial communities.

A microbial community sample (also referred to as the sample) is represented by the hierarchically structured taxa (species, genus, families, etc.) and their relative abundances (also referred to as the community structure), and these species are functioning in concert to maintain stability and adapt to the specific environments (also referred to as the niches or biomes) where the microbial community is living. Samples from the same biome tend to have similar community structures. EBI MGnify contains the most up-to-date biome structure[7] with more than a hundred biomes as of year 2020.

Millions of microbial community samples have already been sequenced and deposited to public databases. It has been noticed that more than 300,000 samples have already been deposited into EBI MGnify database[8], and more have been deposited in NCBI database as of December 2020. However, state-of-the-art methods face difficulties in differentiating these huge amounts of heterogenous samples, making comparison and searching among these samples difficult, while rendering knowledge discovery from samples formidable.

An outstanding objective of microbial community sample search is the identification of samples that are most similar with the query sample, as well as identification of biomes from where the query is most likely to from, namely microbial source tracking. There are already methods that existed for comparison and search of samples. The distance-based methods are the first batches designed for the purpose, whose major strategy is to compare the similarity between two samples. The simplest distance-based method is the Jensen-Shannon Divergence (JSD) measurement[9], which only considered species abundances in the community. More advanced distance-based methods considered both species abundances and their phylogenetic relationships. For example, UniFrac[10] is typical distance-based method, which firstly map their respective sets of taxon abundances on the phylogenetic tree, and secondly traverse the tree and execute operation at each node (each representing a species on the phylogenetic tree) to calculate their similarity. Fast UniFrac[11] and Meta-Storms[12] optimized such procedure by changing tree traversal to array loop. Striped UniFrac[13] further optimized matrix similarity comparison by reorganizing samples. Meta-Prism 1.0[14] uses GPU to accelerate the comparison process. Another batch of methods is based on unsupervised learning, such as SourceTracker[15] based on Bayesian method and FEAST[16] based on Expected-Maximization method. These two unsupervised methods are mostly used for source tracking the biomes from where the samples are most likely to be from. However, current methods are limited by the number of samples against which the comparison and search could be conducted, due to the huge time cost as well as the memory need: Faced with a database with millions of samples from hundreds of biomes, none of the current methods could conduct sample search in a space and time efficient manner, not to say accuracy.

We proposed Meta-Prism 2.0 to solve the large-scale microbial community sample search problem, by means of optimization of data structure and improvement of process flow. (1) It removes redundant nodes to save memory. (2) And it uses a pre-fixed instruction pipeline for speed acceleration. (3) Most importantly, for 1-against-N sample comparison, Meta-Prism 2.0 adopts boost scheme optimization, enabling single instruction multiple data (SIMD) optimization.

Meta-Prism 2.0 has been put to test on several datasets, with largest containing a million of samples. Results have shown that we could achieve at least 20 times speed-up compared to the contemporary approach, such as Meta-Prism 1.0 and Striped UniFrac, when searching against one hundred thousand samples, and Meta-Prism 2.0 is the only method that could handle the search against a million of samples. The memory utilization is also very efficient: at most 20% of memory space is needed by Meta-Prism 2.0 compared to other methods. Though we have saved time and memory by magnitudes, the accuracy is not compromised. For example, Meta-Prism 2.0 obtained 0.99 AUC in microbial source tracking while Striped UniFrac obtained 0.88 AUC on the same dataset. Meta-Prism 2.0 has changed the traditional computational resource-intensive sample search to a cheap and effective procedure that could be conducted by researchers every day, for discovery intricate relationships among samples and mining previously unknown knowledge.

## Methods

Meta-Prism 2.0 has two calculation modes: search mode and matrix mode. The search mode takes two datasets (Query and Target) as input, then outputs each query sample’s top N similar match in target dataset. The matrix mode takes a dataset as input, and outputs a pair-wisely similarity matrix for all samples in the dataset (**Figure 1A**).

**Figure 1.**
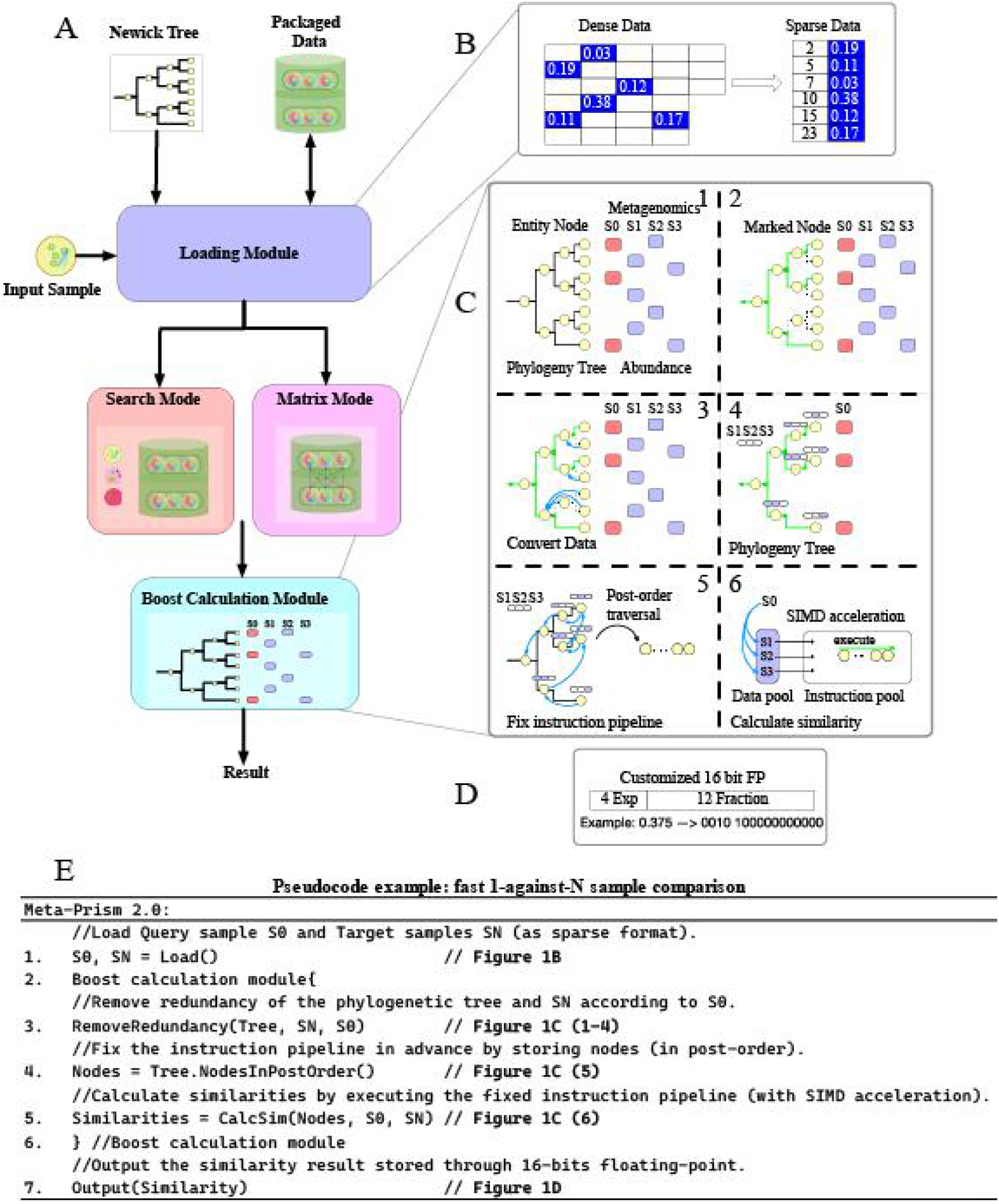
The Meta-Prism 2.0 pipeline with key optimization highlighted. (A) Meta-Prism 2.0 takes taxa abundance as input, maps data to phylogenetic tree, and converts data to sparse format for space optimization. Meta-Prism 2.0 organizes data according to search mode or matrix mode, then sends data to boost calculation module to calculate similarities. (B) Space save scheme packages sample data to sparse format for storage, cut down the usage of both disk and memory. (C) Boost scheme saves resources to the maximum extent by removing redundant nodes without losing sample abundance (1-4), and then fix the instruction pipeline for fast 1-against-N sample comparison (5), followed by SIMD optimization as a compiler-level optimization (6). (D) Similarities are saved in the format of customized 16-bit floating point.

Meta-Prism 2.0 has unlocked several key computational techniques for efficient comparison (**Figure 1**): Firstly, it has utilized a memory-saving data structure (**Figure 1B**). Secondly, to cut down the memory usage, Meta-Prism 2.0 frees memory used by redundant nodes (e.g. nodes represent taxa don’t appear in samples) since these nodes don’t contribute to similarity calculation (**Figure 1C(1-4)**). Thirdly, to further cut down the time usage, Meta-Prism 2.0 fixes the instruction pipeline, discarding redundant instructions before diving into similarity calculation (**Figure 1C(5)**). Fourthly, Meta-Prism 2.0 utilized boost scheme to enable further accelerations through the fixed instruction pipeline and single instruction multiple data (SIMD) optimization (**Figure 1C(6)**). Last but not least, Meta-Prism 2.0 utilizes a customized 16-bit floating-point to store the similarity matrix in a memory-saving manner (**Figure 1D**).

### 2.1 Space-saving data format

For the representation of a single microbial community sample, most of the nodes on the phylogenetic tree are redundant. Current methods use fixed-length array to save sample data with length equals to entities number of phylogenetic tree, wasting considerable amount of memory. Meta-Prism 2.0 store sample data in the form of sparse arrays, saving nodes’ abundances and their indexes in the phylogenetic tree (**Figure 1B**).When calculating similarities, Meta-Prism 2.0 converts sparse data back to dense data with size equals to the taxa number of the phylogenetic tree generated by boost scheme (**Figure 1C(1-4)**). Sparse data structure is applied on disk storage and memory cache, so that the space utilization is reduced on global scale.

The storage scheme is further optimized at the step of similarity result storage. To store similarity results for a sample pair, we designed 16 bits floating-point format with 4 exponential bits and 12 mantissa bits. Considering that similarities are generally between zero and one, we removed two sign bits of exponent and mantissa to increase this format’s arrange and precision (**Figure 1D**).

### 2.2 Boost Scheme for fast 1-against-N sample comparison

We further optimized the time usage and memory usage to the minimum extend through fixing the instruction pipeline and SIMD (Single Instruction Multiple Data). Current methods traverse phylogeny tree (with redundant nodes) and execute operation during similarity calculation, wasting time on redundant operations. To save the time wasted on such redundant operations, we removed redundant nodes before diving into similarity calculation, and fixed the instruction pipeline for each 1-against-N sample comparison, through storing the nodes in post-order, saving time of tree traversal during similarity calculation (**Figure 1C(5)**). We further accelerated the 1-against-N sample comparison utilizing the fixed instruction pipeline and SIMD, so that the time usage could be optimized to the minimum extent (**Figure 1C(6)**).

## Results

### 3.1 Materials

Through manual curation from EBI MGnify database[8], we obtained a dataset consists of 126,274 microbial community samples belonging to 114 different biomes, defined as the Combined dataset. We have also obtained a dataset consists of 10,270 samples belonging to three biomes: Fecal, Human, and Mixed, which have been used in FEAST study[16], defined as the FEAST dataset (**Table 1**). In order to evaluate Meta-Prism 2.0’s speed and memory efficiency on the scale of one million samples, based on extrapolation of MGnify samples, we have also synthesized a dataset with 1,000,010 samples. All samples from these three datasets are accessible from https://github.com/HUST-NingKang-Lab/Meta-Prism-2.0. We used Silva 132 LTPs132 SSU phylogenetic tree[17] in all experiments included in this study.

**Table 1.** The Combined dataset and FEAST dataset used in this study. Details are provided in **Supplementary Table S1**.

| Dataset | Combined dataset | FEAST dataset |
| --- | --- | --- |
| Top-level biome | Root | Human gut |
| Number of biomes involved | 114 | 3 |
| Number of samples | 126,727 | 10,270 |
| Number of species | 45,477 | 5,762 |
| Average number of species per sample | 411.22 | 111.05 |
| Notes | Selected samples from MGnify database | Selected samples from FEAST dataset |

Striped UniFrac, Dynamic Meta-Storms, Meta-Prism 2.0 were compiled by GCC 4.8.5 and ran on CentOS 6.7 with Intel(R) Xeon(R) CPU E5-2678 v3 @ 2.50GHz and 252GB memory. Jensen-Shannon is using Python 3.7.3, SciPy 1.4.1, ran on same CentOS device. The executable Meta-Prism 2.0 steps’ time usage was compiled by clang-1100.0.33.16, and evaluated by Xcode11.5 Instruments Time Profiler, ran on macOS 10.15 with Intel(R) Core(TM) i7-9750H and 32GB memory. Meta-Prism GPU version was compiled by NVCC 10.1, and ran on RTX 2080Ti.

### 3.2 Accuracy evaluation

We evaluated these methods’ accuracies by using simple cross-validation: randomly splitting the dataset into Query dataset and Target dataset, and then searching Query samples (12.5%) against the Target samples (87.5%). We assessed the prediction accuracies, based on checking the consistency of the the predicted biomes and actual biomes of Query samples. For each Query sample, we selected top 100 most similar Target samples, and the contributions of different source biomes of these 100 samples were assessed by SoftMax normalization.

The prediction results were shown in **Figure 2**. On FEAST dataset, every method need to predict testing samples’ biome from 3 biomes: Fecal, Human and Mixed. Distance-based phylogenetic tree approaches (Meta-Prism 2.0, Striped UniFrac and Dynamic Meta-Storms) showed similarly good performance, while Jensen-Shannon Divergence (JSD) obtained lower AUC results. On Combined dataset, every method need to predict testing samples’ biome from 114 biomes. JSD and Dynamic Meta-Storms can’t complete calculation within acceptable time (10 days), so we only compared Meta-Prism 2.0 and Striped UniFrac. Meta-Prism obtained a higher AUC result of 0.9934, while Striped UniFrac’s AUC result was 0.9153.

**Figure 2.**
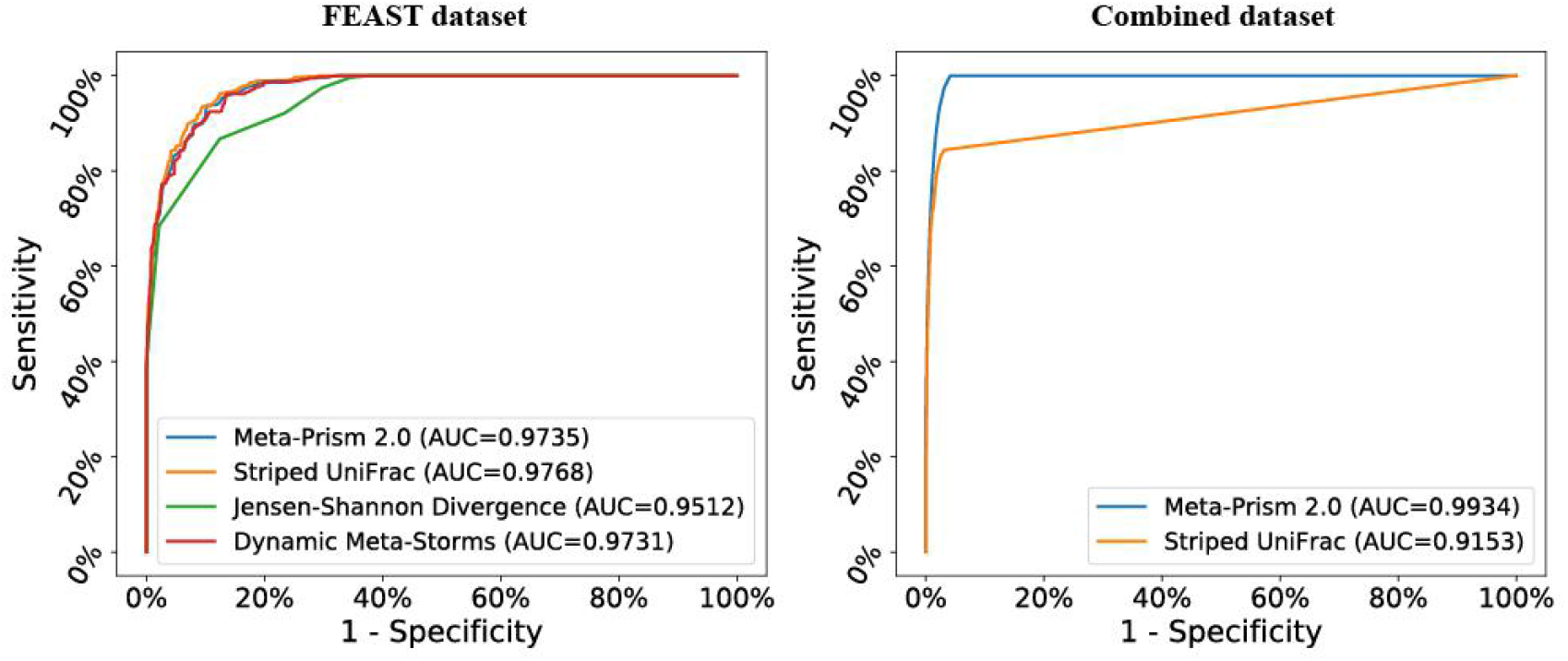
AUC of different methods for sample searches using the FEAST dataset and the Combined dataset.

### 3.3 Computational speed assessment

The time and memory efficiency are the most profound advantage of Meta-Prism 2.0. We first assessed Meta-Prism 2.0’s speed based on using datasets with variate sample sizes, as well as using different number of CPU threads (**Figure 3**). The setting is matrix mode, which takes one dataset as input, then calculates similarities of all sample pairs, and the output is a similarity matrix. The time cost was split into several parts according to computational steps.

**Figure 3.**
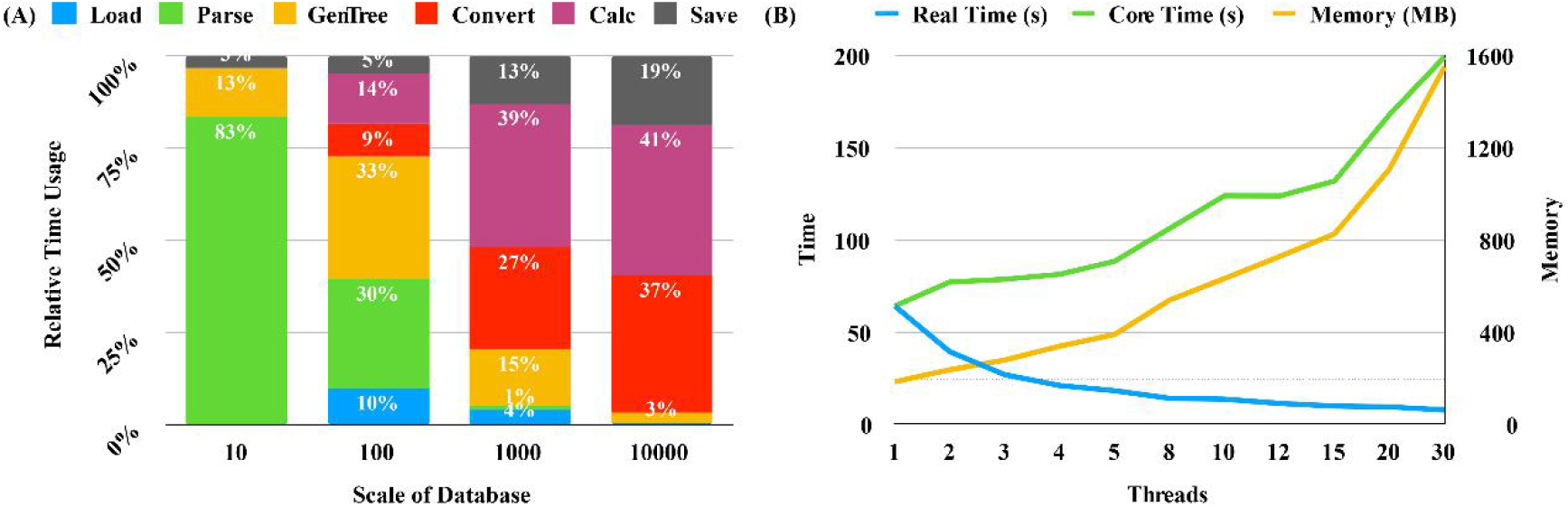
Time usage at different steps and multi-threads performance analysis of Meta-Prism 2.0. (A) Each steps’ time usage with variate sample sizes. Load: load data, Save: save matrix result, Parse: load and parse phylogeny tree, GenTree: generate non-redundant phylogeny tree (without redundant nodes) in boost scheme, Convert: convert sample data from spare format to dense format for sample comparison, Calc: 1-against-N sample comparison. Higher proportion of total time will be used by Convert and Calc steps when the number of sample-pairs increases. (B) Time and memory usage for 10,000 samples’ pair-wise similarity calculation using different number of CPU threads. Real Time: the actual time usage of calculation, Core Time: the sum of each CPU cores’ time usage. Note that since each thread of Meta-Prism 2.0 removes redundancy in the phelogenetic tree and source samples separately, the total memory usage would increase as the number of threads increases.

We also evaluated Meta-Prism 2.0 performance on a dataset with one million samples (see **Materials** for details). Meta-Prism 2.0 can efficiently package one million samples into a 369 MB sized file for storage, and load them within 27 seconds. We transferred the whole workload to a laptop and search 100 samples against this dataset with a single CPU thread. It costed 324.96 seconds (less than 6 minutes) CPU time to complete the search using only 6.9 GB memory. So far as we know, Meta-Prism 2.0 is the only the only method that could handle the search against a million of samples.

### 3.4 Computational speed comparison

We further selected different sizes of samples from the Combined dataset (10, 100, 1,000, 10,000, 100,000 and 126,727) to compared different methods. The setting is again matrix mode. We compared time and memory usage of Striped UniFrac, Dynamic Meta Storms, JSD, Meta-Prism GPU and Meta-Prism 2.0. Meta-Prism GPU is the only method that uses GPU for calculation, and we take real time usage as its time usage. While we take CPU core time usage as other methods’ time usage. JSD and Meta-Storms can’t calculate similarity matrix when samples’ size is or higher than 10K within acceptable time (10 days).

Results have shown that Meta-Prism 2.0 could achieve superior performance on both time usage and memory usage (**Figure 4)**. Specifically, when the sample size is no more than one thousand, Meta-Prism 2.0 used similar core time compared with Striped UniFrac **(Figure 4A)**. When dataset size became larger, the gap between Meta-Prism 2.0 and Striped UniFrac became larger. When calculating similarity matrix for Combined dataset (generating 126,727*126,727 similarity matrix), Meta-Prism 2.0 is 55 times faster than Striped UniFrac. Meta-Prism GPU’s real time usage is smaller than Meta Prism 2.0 core time usage. However, when Meta-Prism 2.0 uses 3 CPU cores or more it will be faster than Meta-Prism GPU.

**Figure 4.**
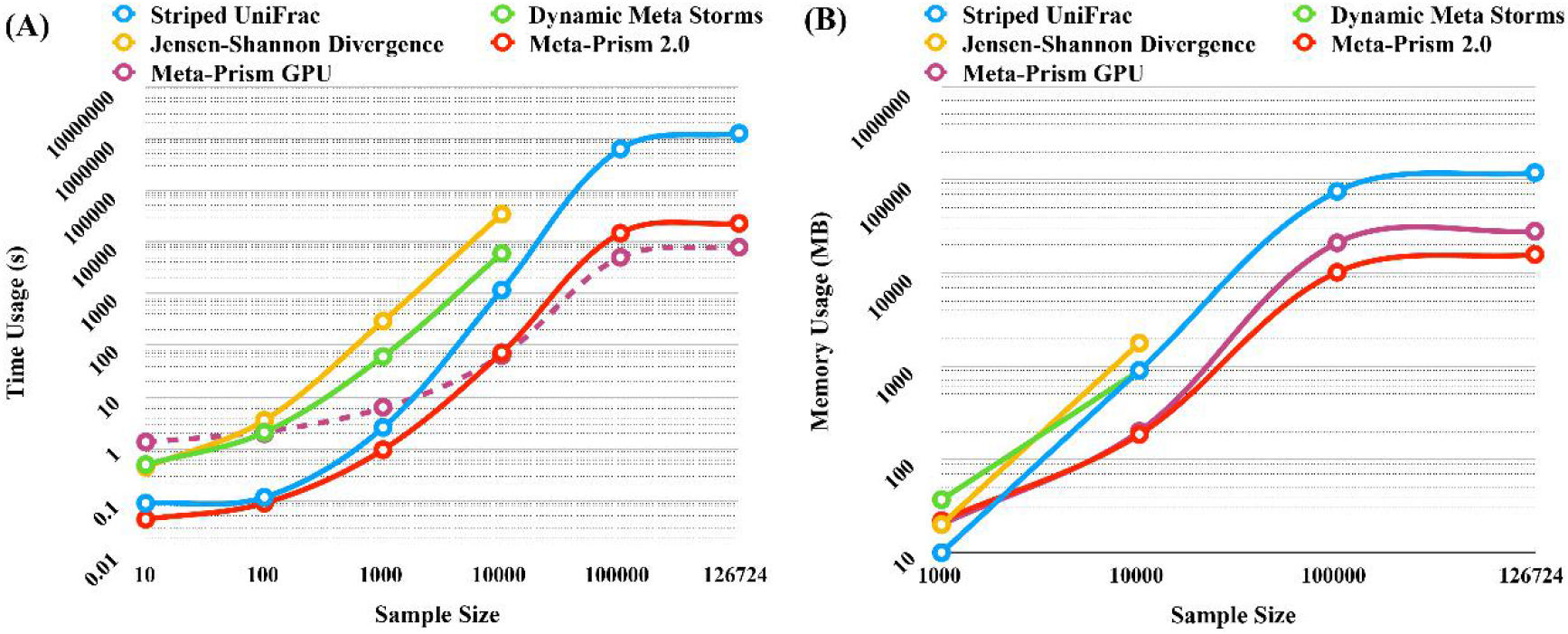
Time and memory usage of samples when calculating similarity matrix for datasets with different number of samples. (A) is for time usage comparison, and (B) is for memory usage comparison. In (A), Meta-Prism GPU time usage with dash line is GPU time usage, others are CPU core time usage.

When the dataset size is smaller than 1,000, all methods utilized similar memory space. However, Meta-Prism 2.0’s memory usage was only 11.1% of Striped UniFrac’s when calculating similarity matrix for Combined dataset with more than 100,000 samples. The utilization of customized 16 bits floating point was the key reason behind such efficient memory use: as the memory occupied by the similarity matrix increases quadratically when the dataset size increases, the little saving of a sample-pair’s similarity storage would lead to amplified reduction of memory usage.

Since Meta-Prism 2.0 is ultrafast, it is natural to wonder how far is the speed of Meta-Prism 2.0 to the theoretical lower bound for sample search. To answer this question, we took IO Only as the lower bound for sample search, in which we only record time used for loading data and writing matrix calculation result (**Figure 5**). The result have shown that Meta-Prism 2.0 is already close to the limit of optimization: on datasets of different sizes, the time costs of Meta-Prism 2.0 were only 2 times of IO Only, while magnitude smaller than those of Striped UniFrac.

**Figure 5.**
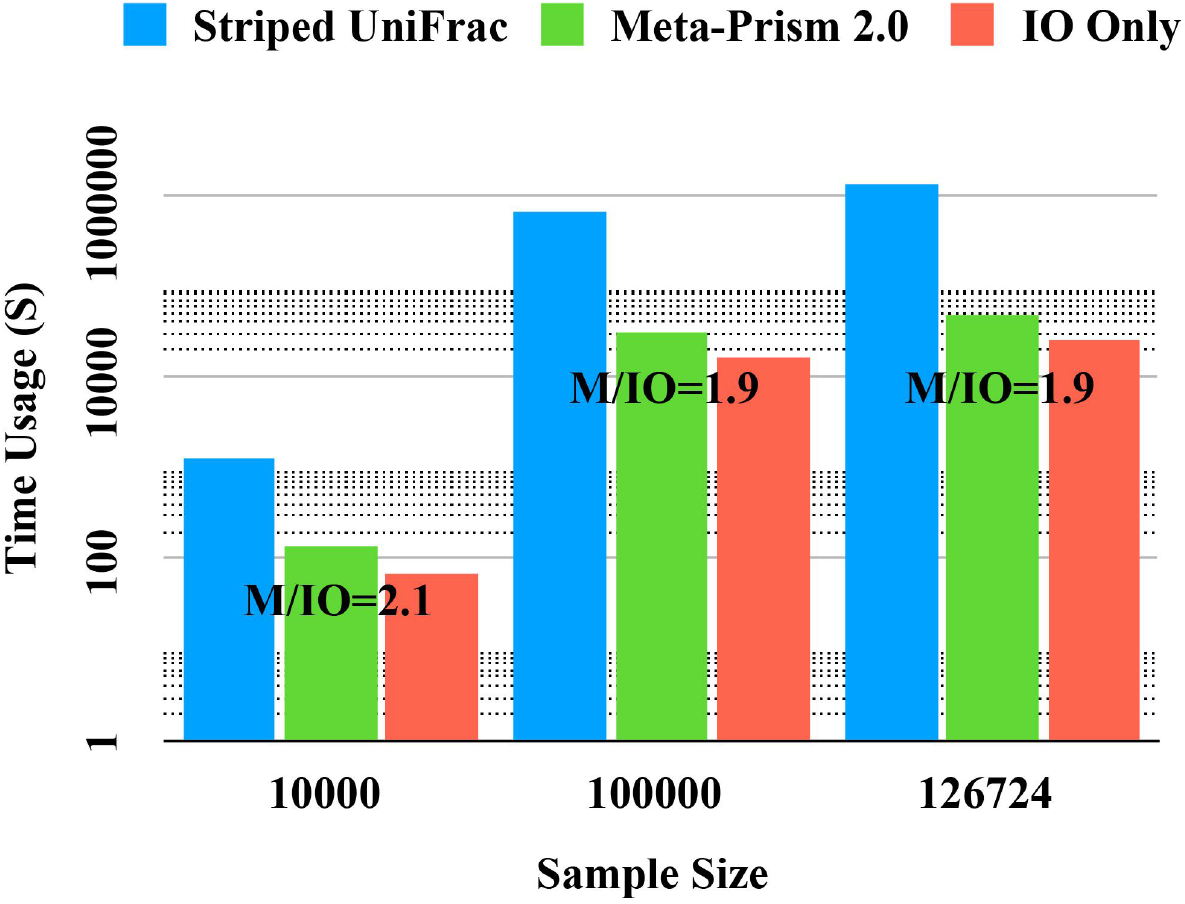
Time usage for different methods and IO Only on datasets with different sizes. “M/IO” is the ratio of time cost of Meta-Prism 2.0 over that of IO Only.

### 3.5 Real data applications

Meta-Prism 2.0 is able to precisely identify the biome for samples of unknown origin, thus enabling the source tracking of samples. For example, it enables accurate differentiation of samples from close biomes such as “human skin” and “human oral” (the first application), identification of the biome for samples with unclear origin (the second application), as well as detection of microbial contamination (the third application).

Firstly, we have tested Meta-Prism 2.0’s ability in accurate differentiating of samples from close biomes. We have obtained 1,261 skin metagenomic samples (MGYS00005172)[18] and 70 oral metagenomic samples (MGYS00005569)[19] from MGnify[8]. We used Meta-Prism 2.0 to calculate the similarities matrix of 1,331 samples on a laptop, which cost only 3.75 seconds and 11MB memory. Then we clustered samples by their similarities, by using affinity propagation from Scikit-learn 0.20.3. The samples were successfully clustered into two groups whose sizes are 1,260 and 71 (**Figure 6**). Within total of 1,331 samples, only 9 samples (5 skin samples and 4 oral samples) were miss-clustered, proving Meta-Prism 2.0’s ability of fast and accurate differentiation of samples from close biomes.

**Figure 6.**
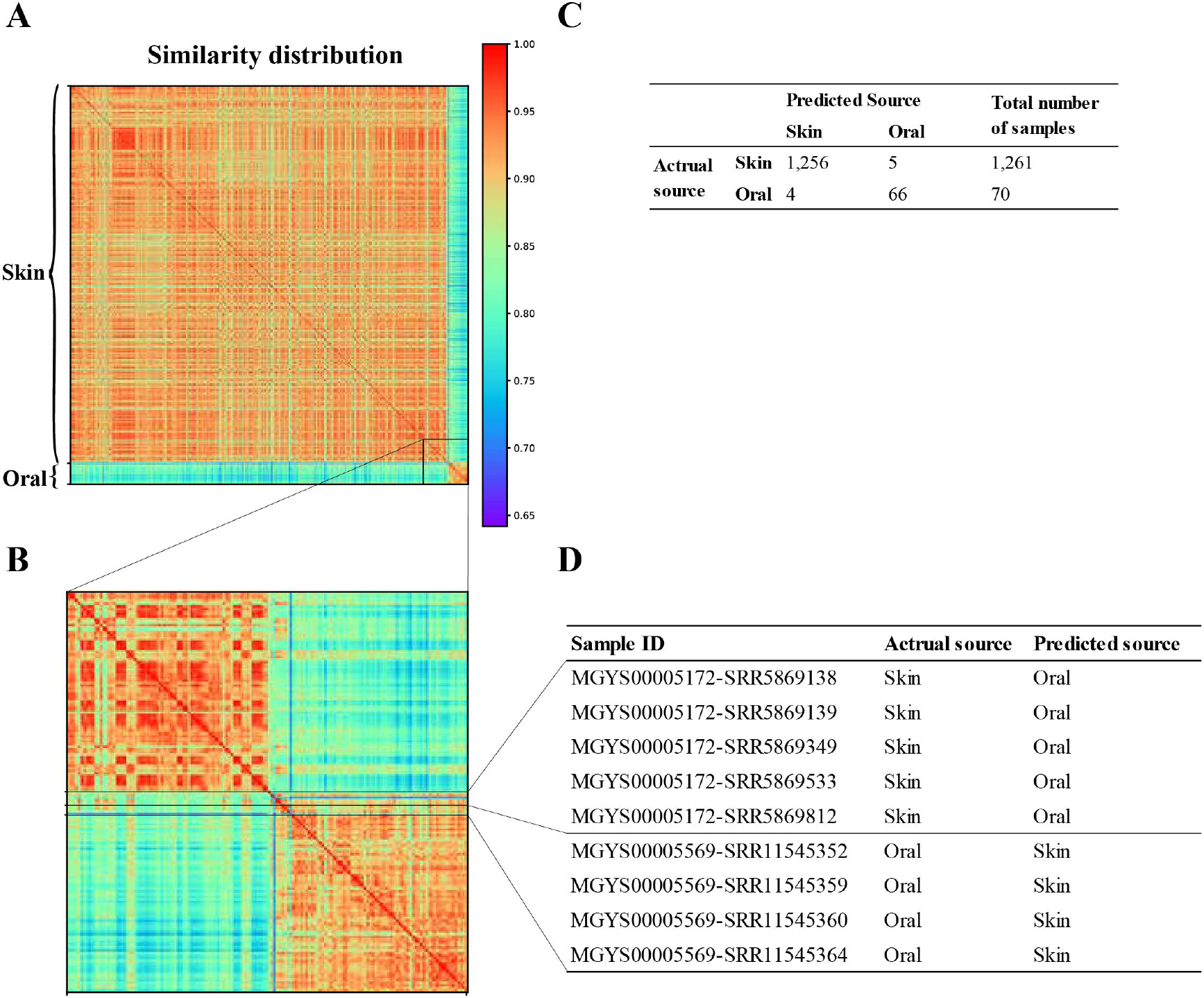
Cluster result of human samples from close biomes using similarities calculated by Meta-Prism 2.0. (A) Similarity distribution of 1,331 samples. The samples were successfully clustered into two groups though we didn’t give a certain number of clusters. (B) Similarity distribution of 9 mis-clustered samples. (C) Confusion matrix and the number of samples with in each actual source biome and predicted biome. (D) MGnify study accession, run accession, actual biome source and predicted biome source of 9 mis-clustered samples.

Secondly, we have evaluated the performance of Meta-Prism 2.0 on source tracking environmental samples from less studies biomes, based on searching 11 groundwater samples curated from Saudi Arabian (MGYS00001601)[20] against the combined dataset. The biome “groundwater” is less studied, with a handful of samples in the combined dataset (MGYS00005245). Results have shown that Meta-Prism 2.0 could successfully identify source-related biomes for samples from “groundwater”. Within top 100 most similar community samples for each “groundwater” query sample, there are on average 64 groundwater-related samples for each Query sample. (“root-Environmental-Terrestrial”, “root-Environmental-Aquatic”, “root-Engineered-Wastewater” and “root-Host-associated-Plants”). Nevertheless, there is no “groundwater” sample in the top 100 similar samples searched by Meta-Prism 2.0. This could largely due to the fact that, “groundwater” samples in the combined dataset that we used are curated from New Zealand, which are in nature drastically different from our Query samples.

Finally, we have evaluated the Meta-Prism 2.0’s power in detecting contaminations. We investigated the contamination of indoor house surfaces community, by selecting 611 samples from indoor house surfaces in Chicago as Query samples and searching against 6,285 samples (899+3,773+721+692 from “human skin”, “environmental”, “mammal”, and “plants”, respectively). The analysis cost only 6.16 seconds to complete. Our results shown that most closed biome source for indoor house surfaces samples is “human skin” (average similarity 0.889), indicating that there is a large proportion of microbial community contamination from human skin. This agrees with previous analyses by SourceTracker[15] and FEAST[16]. Again, it has proved the ability of Meta-Prism 2.0 for accurate and fast microbial community contamination screening.

## Discussions and Conclusion

In this work, we have designed Meta-Prism 2.0 as an ultrafast and memory efficient approach to search microbial community samples against millions of samples. The sample search problem has encountered great difficulties when faced with millions of samples to be searched against, largely due to the computational space and time limitations. These issues should be solved or optimized to cope with the search against millions of samples. Meta-Prism 2.0 has been designed based on the sparse data structure, time-saving tree traversal and to-do queue techniques, thus it has enabled ultra-fast, accurate and memory-efficient search among millions of samples.

Results have shown that compared to the current methods serving the same purpose, Meta-Prism 2.0 is at least 20 times faster, while memory efficiency is at least 4 times smaller. Additionally, the speed of Meta-Prism 2.0 is already very close to lower bound of the search, leaving little room for further improvement for distance-based methods. Furthermore, according to our experiment, Meta-Prism 2.0 can even store the community structure of all samples from EBI MGnify dataset (300,000 in total as of Oct. 2020) on a laptop and searching against it at an acceptable speed. Finally, we have provided several concrete examples, which have proven the effectiveness of Meta-Prism 2.0 in knowledge discovery.

In summary, Meta-Prism 2.0 can perform searches among millions of samples with very low memory cost and fast speed, enabling source tracking and knowledge discovery from samples mining at a massive scale. Meta-Prism 2.0 has changed the traditional resource-intensive sample comparison and search to a cheap and effective procedure that could be conducted by researchers everyday, for mining intricate relationships among samples as well as for discovery of previously unknown knowledge.

## Acknowledgments

This work was partially supported by Ministry of Science and Technology’s grant 2018YFC0910502, National Undergraduate Training Program for Innovation and Entrepreneurship of China (Program No. 201910487071), and National Science Foundation of China grant 31871334 and 31671374.

